# Assessing Runs of Homozygosity: A comparison of SNP Array and Whole Genome Sequence low coverage data

**DOI:** 10.1101/160705

**Authors:** Francisco C. Ceballos, Scott Hazelhurst, Michèle Ramsay

**Affiliations:** Sydney Brenner Institute for Molecular Bioscience, Faculty of Health Sciences, University of the Witwatersrand, Johannesburg, South Africa.; Division of Human Genetics, School of Pathology, Faculty of Health Sciences, University of the Witwatersrand, Johannesburg, South Africa.; School of Electrical & Information Engineering, University of the Witwatersrand, Johannesburg, South Africa.

**Keywords:** Runs of Homozygosity, ROH, SNP array data, WGS low coverage data

## Abstract

Runs of Homozygosity (ROH) are sequences that arise when identical haplotypes are inherited from each parent. Since their first detection due to technological advances in the late 1990s, ROHs have been shedding light on human population history and deciphering the genetic basis of monogenic and complex traits and diseases. ROH studies have predominantly exploited SNP array data, but are gradually moving to whole genome sequence (WGS) data as it becomes available. WGS data, covering more genetic variability, can add value to ROH studies, but require additional considerations during analysis. Using SNP array and low coverage WGS data from 1885 individuals from 20 world populations, our aims were to compare ROH from the two datasets and to establish software conditions to get comparable results, thus providing guidelines for combining disparate datasets in joint ROH analyses. Using the PLINK Homozygosity functions, we found that by allowing 3 heterozygous SNPs per window when dealing with WGS low coverage data, it is possible to establish meaningful comparisons between data using the two technologies.

## Introduction

Runs of Homozygosity (ROH) are contiguous regions of the genome where an individual is homozygous across all sites. ROH arise when two copies of an ancestral haplotype are brought together in an individual. Consequently, that haplotype would be autozygous, *i.e.* homozygous by descent. ROH were first discovered using genome-wide microsatellite scans in the mid 1990s (Broman et al. 1999). Members of two families recruited to construct the first human genetic maps carried 4-16 ROH typically 2-40 cM in length; the most extreme individuals had a total of 253 cM in ROH, consistent with close inbreeding. Henceforth, ROH were found to be ubiquitous even in outbred populations; indeed, we are all inbred to some degree and ROH captures this aspect of our individual demographic histories (Biraben 1980; Donnelly 1983; Gibson et al. 2006; Keller et al. 2011).

ROHs have been a subject of study since their discovery due to their applications for understanding human population and disease genetics (Szpiech et al. 2013; Joshi et al. 2015). The number and length of ROH reflect individual and population history while the homozygosity burden can be used to investigate the genetic architecture of complex disease (McQuillan et al. 2008; Pemberton et al. 2012; Ghani et al. 2015; Yang et al. 2015; Howrigan et al. 2016; Johnson et al. 2016). In view of their usefulness the number of articles published using ROH has been increasing significantly in the last years (162 in 2005, 322 in 2010 and 620 in 2016, PubMed search using R package RISmed) using predominantly DNA SNP array genotypes. It is expected that, with the current availability of full genome sequences, ROH will be used extensively as an augmentative approach to study population structure, demographic history and in deciphering the genetic structure of complex diseases.

The first aim of this article is, therefore, to compare the outcomes and general conclusions drawn for array-based data and low coverage (3-6x) whole genome sequence data from the same groups of individuals. The second is to obtain appropriate parameters of ROH calling that allow meaningful comparison between ROH obtained from both technologies.

There are two major methods for identifying ROH: observational genotype-counting algorithms (Purcell et al. 2007) and model based algorithms (Pemberton et al. 2012). Observational approaches use algorithms that scan each chromosome by moving a fixed size window along the whole length of the genome in search of stretches of consecutive homozygous SNPs (Purcell et al. 2007). This approach is implemented in PLINK v1.9 where a given SNP is considered to potentially be in an ROH by calculating the proportion of completely homozygous windows that encompass that SNP. If this proportion is higher than a defined threshold, the SNP is designated as being in a ROH. In the algorithm, a variable number of heterozygote positions or missing SNPs can be specified per window in order to tolerate genotyping errors and failures. An ROH is called if the number of consecutive SNPs in a homozygous segment exceeds a predefined threshold in terms of SNP number and/or covered chromosomal length. The simplicity of the approach used by PLINK allows efficient execution on data from large consortia (Joshi et al. 2015). On the other hand, haplotype-matching algorithms (e.g. Germline) (Gusev et al. 2009) for calculation of identity-by-descent (IBD) can also be used to identify ROH, as a special case of IBD within an individual. Model-based approaches use Hidden Markov Models (HMM) to account for background levels of LD, like the one implemented in Beagle (Browning and Browning 2010). Tests on simulated and real data showed that the approach using PLINK outperformed Germline and Beagle in detecting ROH (Howrigan et al. 2011). These models have been used to find ROH in array and WGS data; however, the HMM model approach is also used with Whole Exome Sequence (WES) data as an alternative to discover SNP variants and small to medium length ROH (Zhuang et al. 2012; Mezzavilla et al. 2015). However, with the sparse nature of the WES target design, long ROH detection is not possible. Specific software, like “homozygosity heterogeneous hidden Markov model (HMM)” or H3M2, was designed to deal with this type of data (Magi et al. 2014).

Accurate ROH calling requires high density SNP genome-wide scan data. A number of factors influence the quality of ROH calling, including the marker density, their distribution across the genome, the quality of the genotype calling/error rates and minor allele frequency. Currently ROH studies have been carried out using genome-wide scan data overwhelmingly from SNP arrays (Ku et al. 2011; Yang et al. 2012; Joshi et al. 2015), both because of the availability of this data and the fact that array data is considered the gold standard with very low genotyping calling error rates (typically <0.001). However SNP arrays usually include ∼1-2.5 million SNP typically with allele frequencies >0.05, chosen to best represent haplotype structure in target populations. Arrays with more than 300k SNP genome-wide coverage have been shown to be good enough to successfully detect ROH longer than 1 Mb, which correspond to true ROH arisen by autozygosity (McQuillan et al. 2008). Indeed, it is expected that long ROH will keep their homozygous status independently of the SNP coverage. However, the relative sparsity of SNPs on an array may mean that true heterozygous SNPs between the markers on the array may be missed, thereby making two close-by ROHs appear as one, longer ROH.

A WGS approach, on the other hand, assays every variant so all accessible bases can now be genotyped and more than several million variants, from the most common to the most private can be obtained for each individual (Nielsen et al. 2011; Goodwin et al. 2016). For cost reasons, low coverage sequencing is often employed to maximize the number of participants in a study and strengthen its power. In this case rare SNPs are called significantly less often, with higher error rates, than common SNPs. Whole genome sequence with low-coverage (e.g. 4x average) has a high probability that only one of the two chromosomes of a diploid individual has been sampled at a specific site (Nielsen et al. 2011; Goodwin et al. 2016). Error rates of low coverage WGS can get up to 15% or higher. Of course, reducing and quantifying the uncertainty associated with SNP calling may be accomplished using sophisticated algorithms, and this approach has been subject to extensive research (Goodwin et al. 2016). However, the error rate for low coverage WGS is significantly higher than for array data, which will lead to inaccuracy in ROH calling. This is particularly important, as the cost of WGS becomes more affordable and data more available (Wetterstrand 2016), opening up new possibilities to study ROH in greater detail, replicate results from SNP array data studies, or to the study the relationship of ROH, especially shorter ones, with new populations or traits. Hence, parameters of ROH calling algorithms require tuning to the characteristics of the underlying data in order to obtain meaningful comparable results between studies using different technologies. While in the long run, high coverage data (>30x) will become more affordable, for the medium-term at least, low-coverage WGS data will be an important source for many analyses.

## Results

### Comparing variant calling between technologies

In order to have a meaningful comparison of ROH obtained from array and WGS low coverage data it is important to first analyze the differences in presence of heterozygous SNPs and variant calling between both technologies. To assess the error rate in heterozygote calling in the WGS, the percentage of concordance in the variant calling between the array and the WGS data, is shown for every population studied (Table 1). As expected, WGS included more heterozygotes SNPs since the SNP array captured only data from ∼2.5M nucleotide positions in the autosomal genome, whereas the WGS provided data for the entire length of the genome (∼2.8×10^9^ nucleotide positions). On average, for all the populations analyzed, the WGS low coverage data had 6.3 times more heterozygous SNPs (2,558,000 ±71,700) compared to the array (404,700 ±7,717) (Table 1). In WGS data there is 1 heterozygous SNP per 1.1 Kb vs 1 in 7.1 Kb in array data. On average the concordance in variant calling by array and WGS is 99.6% (±0.05%). Of the 0.4% (±0.05) discordant calling, on average, 0.1% (±0.03) of the SNPs was called heterozygous by the array and homozygous by WGS and 0.3% (±0.02) of the SNPs was called heterozygous by WGS, but homozygous by array. Considering that array genotyping is the gold standard, WGS data, on average, led to erroneous calling of 0.3% (±0.02) of heterozygous SNP, which would incorrectly be reported as a break in a given ROH. On average, for all the populations, there will be 6,500 SNPs (±714) per individual wrongly called as heterozygous, and that is roughly 2.4 SNP (±0.3) per Mb. This error rate is however different across the studied populations, with the JPT having the higher error rate (13,000 wrongly called heterozygotes; 4.5 SNPs per Mb) and the ZUL having the lowest (740 wrongly called heterozygotes; 0.3 SNPs per Mb).

**Table 1.** Mean number of heterozygote SNPs (per called SNP) in array and WGS low coverage data for 20 world populations

|  | VARIANT CALLING |  |  |  |  |  | ROH |
| --- | --- | --- | --- | --- | --- | --- | --- |
|  | Ave N of<br>Het. WGS | Ave N of<br>Het. Array | Concor. | Discor. | He A – Ho W | Ho A – He W | error |
| FIN | 2 432 921.7 | 398 280.1 | 99.6929 | 0.3071 | 0.0402 | 0.2669 | 0.227 |
| GBR | 2 463 526.4 | 405 223.1 | 99.6898 | 0.3102 | 0.0430 | 0.2672 | 0.224 |
| IBS | 2 440 125.2 | 399 870.1 | 99.6547 | 0.3453 | 0.0412 | 0.3041 | 0.263 |
| TSI | 2 445 524.4 | 401 124.4 | 99.6015 | 0.3985 | 0.0424 | 0.3562 | 0.314 |
| CEU | 2 479 523.5 | 417 837.2 | 99.6365 | 0.3635 | 0.0402 | 0.3232 | 0.283 |
| ACB | 3 283 726.5 | 454 173.7 | 99.6723 | 0.3277 | 0.0403 | 0.2874 | 0.247 |
| ASW | 3 262 716.1 | 462 107.3 | 99.6526 | 0.3474 | 0.0448 | 0.3026 | 0.258 |
| MXL | 2 524 698.2 | 385 362.1 | 99.7197 | 0.2803 | 0.0433 | 0.2370 | 0.194 |
| CLM | 2 317 649.7 | 377 844.5 | 99.6716 | 0.3284 | 0.0460 | 0.2825 | 0.236 |
| PEL | 2 100 245.2 | 352 485.3 | 99.6987 | 0.3013 | 0.0411 | 0.2601 | 0.219 |
| PUR | 2 421 174.0 | 381 603.3 | 99.4125 | 0.5875 | 0.0448 | 0.5427 | 0.498 |
| CDX | 2 313 375.1 | 371 361.9 | 99.7197 | 0.2803 | 0.0351 | 0.2452 | 0.210 |
| CHB | 2 330 226.6 | 377 553.5 | 99.7197 | 0.2803 | 0.0366 | 0.2437 | 0.207 |
| CHS | 2 317 649.7 | 377 844.5 | 99.6991 | 0.3009 | 0.0451 | 0.2558 | 0.211 |
| JPT | 2 320 417.1 | 375 586.6 | 99.3659 | 0.6341 | 0.0354 | 0.5988 | 0.563 |
| KHV | 2 350 584.8 | 368 521.5 | 99.8549 | 0.1451 | 0.0343 | 0.1109 | 0.077 |
| YRI | 2 840 113.4 | 463 890.4 | 99.5746 | 0.4254 | 0.0383 | 0.3871 | 0.349 |
| LWK | 2 840 253.9 | 441 435.7 | 99.5545 | 0.4455 | 0.0457 | 0.3998 | 0.354 |
| ZUL | 2 840 578.1 | 441 536.6 | 98.7062 | 1.2938 | 0.6338 | 0.6600 | 0.026 |
| BAG | 2 840 658.6 | 441 412.3 | 99.2791 | 0.7209 | 0.3157 | 0.4052 | 0.090 |

Mean variant calling concordance (in %) is shown. Discordance is decomposed in SNP called heterozygous by array but homozygous by WGS and vice versa. Finally a ROH error is defined as the % of SNP that according to variant calling discordance would break ROH in WGS low coverage data.
**Concor.** Concordant, **Discor.** Discordant
**He A – Ho W** SNP called heterozygote by array and homozygote by WGS
**Ho A – He W** SNP called homozygote by array and heterozygote by WGS
**ROH error** % of SNPs that being wrongly called can break a ROH

### Assessing the impact PLINK tolerating heterozygous SNPs in the search for ROH

PLINK software, by allowing a certain number of heterozygous SNPs per window (the default value being 1 heterozygous SNP per window), already takes into account possible calling errors that may wrongly break a long ROH. By allowing this heterozygous SNP, the software produces an error that depends on the number of SNP (in homozygous state) per ROH. Figure 1 shows *ep(P,h)*, a measurement of the empirically observed number of heterozygous SNPs found in ROHs in population *P* when allowing *h* heterozygous SNPs (1 heterozygous SNP in the array data and 1 to 5 in WGS data). This figure shows that for most of the populations the *ep(P,h)* produced by allowing a single heterozygous SNP per ROH in array data is equivalent to allowing 4 to 5 heterozygous SNPs in WGS data. A few populations deviated from this observation: TSI (0.27% for the array data vs 0.17% after allowing for 5 heterozygous in WGS data), ASW (0.185 vs 0.122), ACB (0.185 vs 0.138), YRI (0.13 vs 0.114), BAG (0.161 vs 0.133) and ZUL (0.136 vs 0.105). These differences are provoked by differences in the mean number of SNPs per ROH as it can be seen in Table 1 of the Supplementary material (Supplemental_Table_S1). For example, the TSI population has, on average, 368 SNPs in the homozygous state per ROH in the array data, less than half of the average SNP per ROH in array data across all populations (714.7).

**Figure 1.**
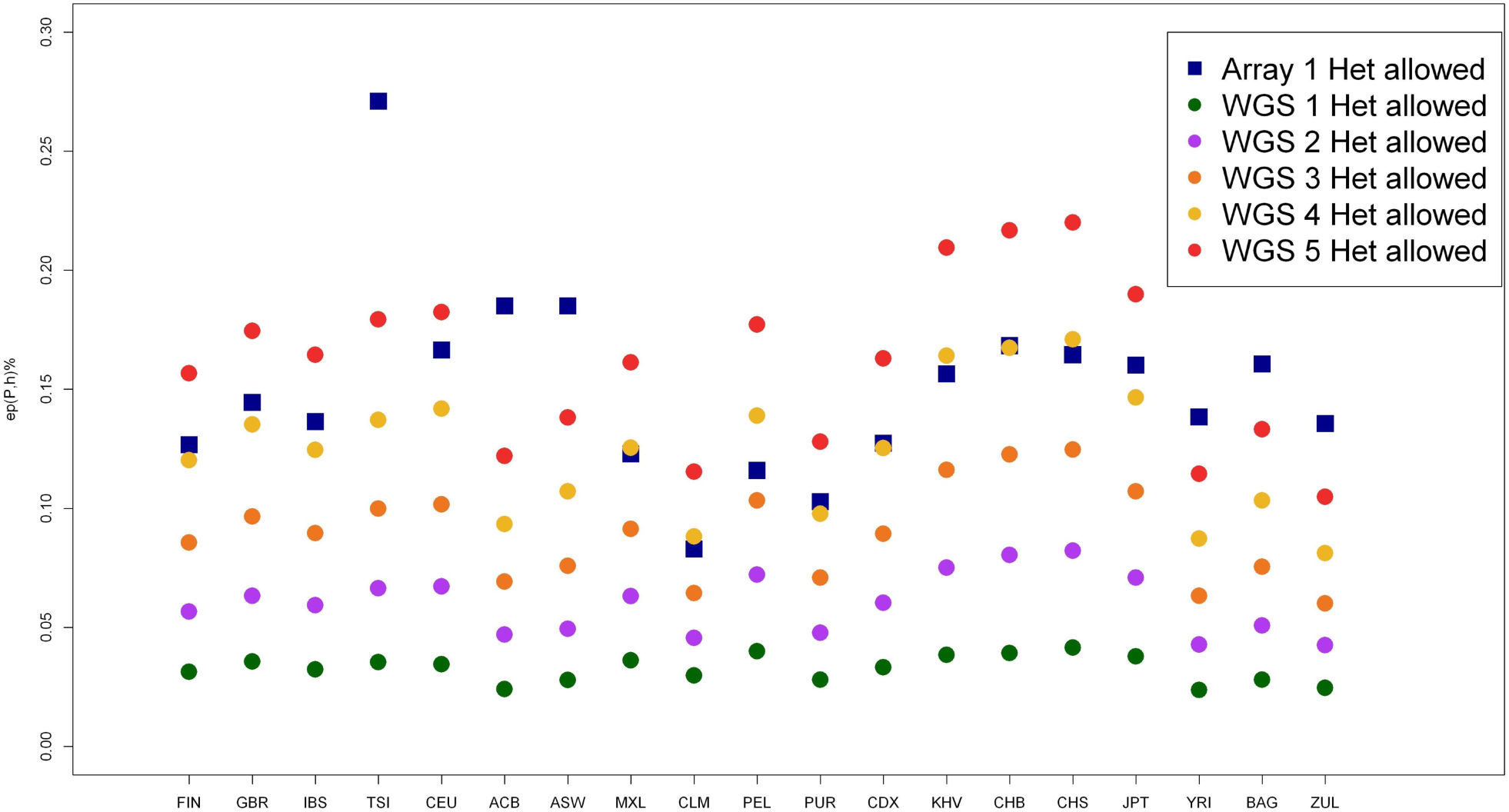
Effect of allowing heterozygous SNPs per window evaluated by *ep(P,h)* as a measure of the empirically observed actual number of heterozygous SNPs found in ROHs in population *P* when we allow *h* heterozygous SNP. (See supplementary material for the definition).

### Obtaining equivalent ROH estimates using data from both technologies

According to both Table 1 and Figure 1 it seems appropriate to compare ROH from both technologies allowing 1 to 5 heterozygote SNPs in WGS data in order to obtain equivalent results. Violin plots show the distribution of mean number of ROH (Figure 2), mean ROH size (Figure 3) and mean total sum of ROH (Figure 4) per population and using array data, compare to WGS data with 1, 2 or 3 tolerated heterozygotes per window. Almost without exception, the distribution between array and WGS data is most similar when 3 heterozygous SNPs in the WGS data are allowed per window. Mean values and standard deviations for up to 5 heterozygous SNPs allowed per window are shown in Supplementary Table 2 (Supplemental_Table_S2). Figure 5 (top row of images) shows the correlations with the array data as heat-maps between number of ROH (i), mean ROH size (ii), and total sum of ROH (iii) for each population and a different number of allowed heterozygous SNPs in the WGS data (values and probabilities shown in Supplemental_Table_S3). The correlations, as expected, increase with more heterozygous SNPs being allowed in the WGS data. Correlations are not homogeneous, south and East Asian populations show lower correlations in comparison with other populations. An alternative representation by line charts is shown in Supplementary Figure 1 (Supplemental_Fig_S1), where differences between populations are perceived more easily. Results of the statistical comparison between ROH obtained from array and WGS (with a different number of heterozygous SNPs allowed) (Figure 5 bottom row of images) by the Mann-Whitney-Wilcoxon (MWW) nonparametrical test are shown as a heat-map of significance (p values; blue = not significant) in Figure 5 bottom row of images. P-values are presented in Supplementary Table 3 (Supplemental_Table_S3). Figure 5 shows heterogeneous results across populations. In general, by allowing 3 heterozygotes SNPs per window in WGS the statistical outcomes in the number of ROH, mean ROH size and total sun of ROH are similar between array and WGS data. However, Figure 5 bottom row also shows that for the Asian populations, especially the JPT, for the number of ROH and total sum of ROH differences between array and WGS data are significant for every heterozygous SNP allowed.

**Figure 2.**
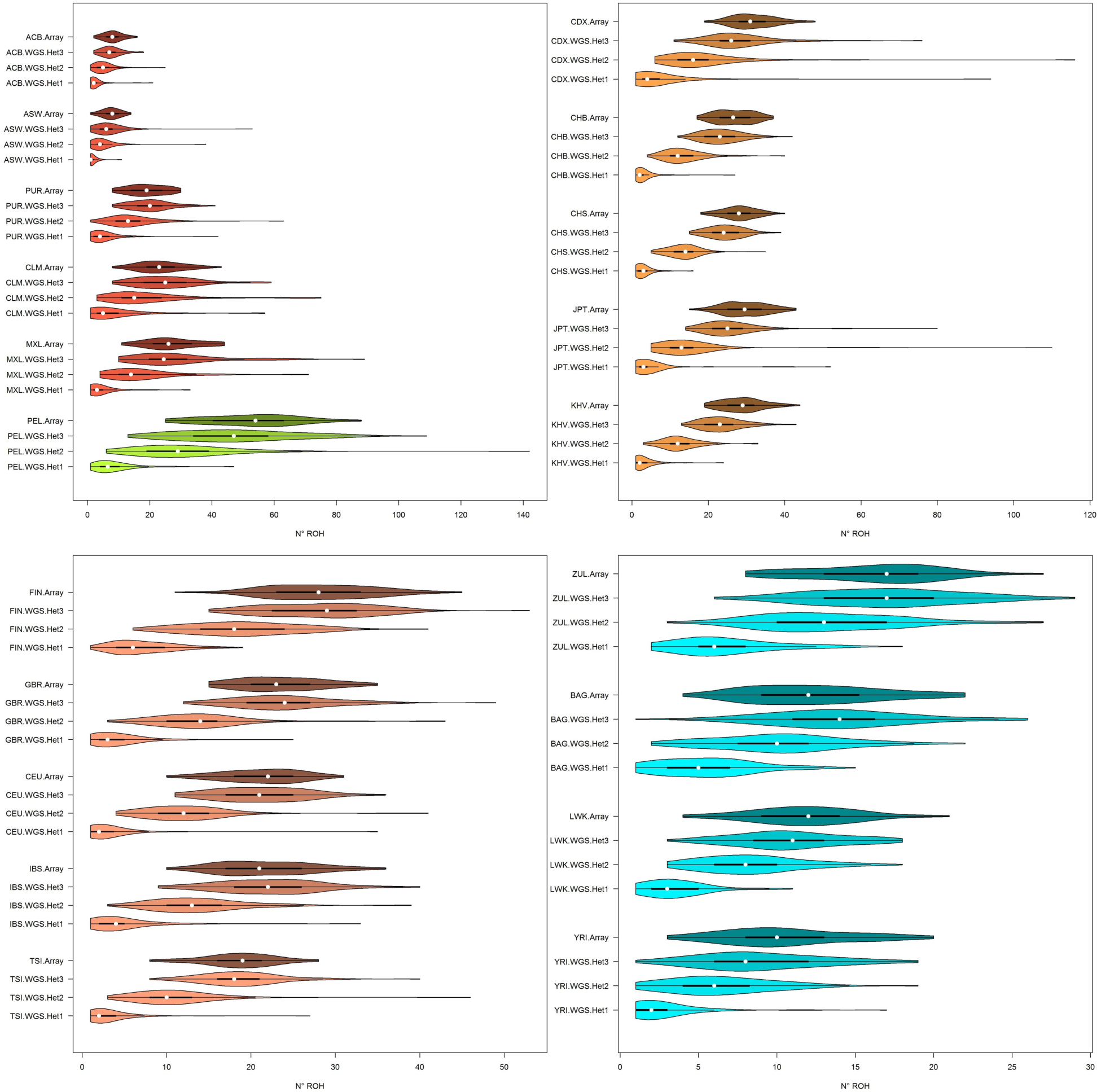
Violin plots of the mean number of ROH longer than 1 Mb. Populations are colored by 5 biogeographical groups by admixture analysis. Admixed (Hispanic-American: CLM, MXL; African-American: ACB, ASW) – blue, Native Americans (PEL) – green, East (CHS, CDX, JPT) and South (KHV) Asia – tan, North (FIN, GBR, CEU) and South (IBS, TSI) Europe – violet, South (ZUL), East (BAG, LWK) and West (YRI) Africa – red. Four distributions per population are shown, array data with 1 heterozygous SNP allowed per window and WGS with 1 to 3 heterozygous SNPs allowed.

**Figure 3.**
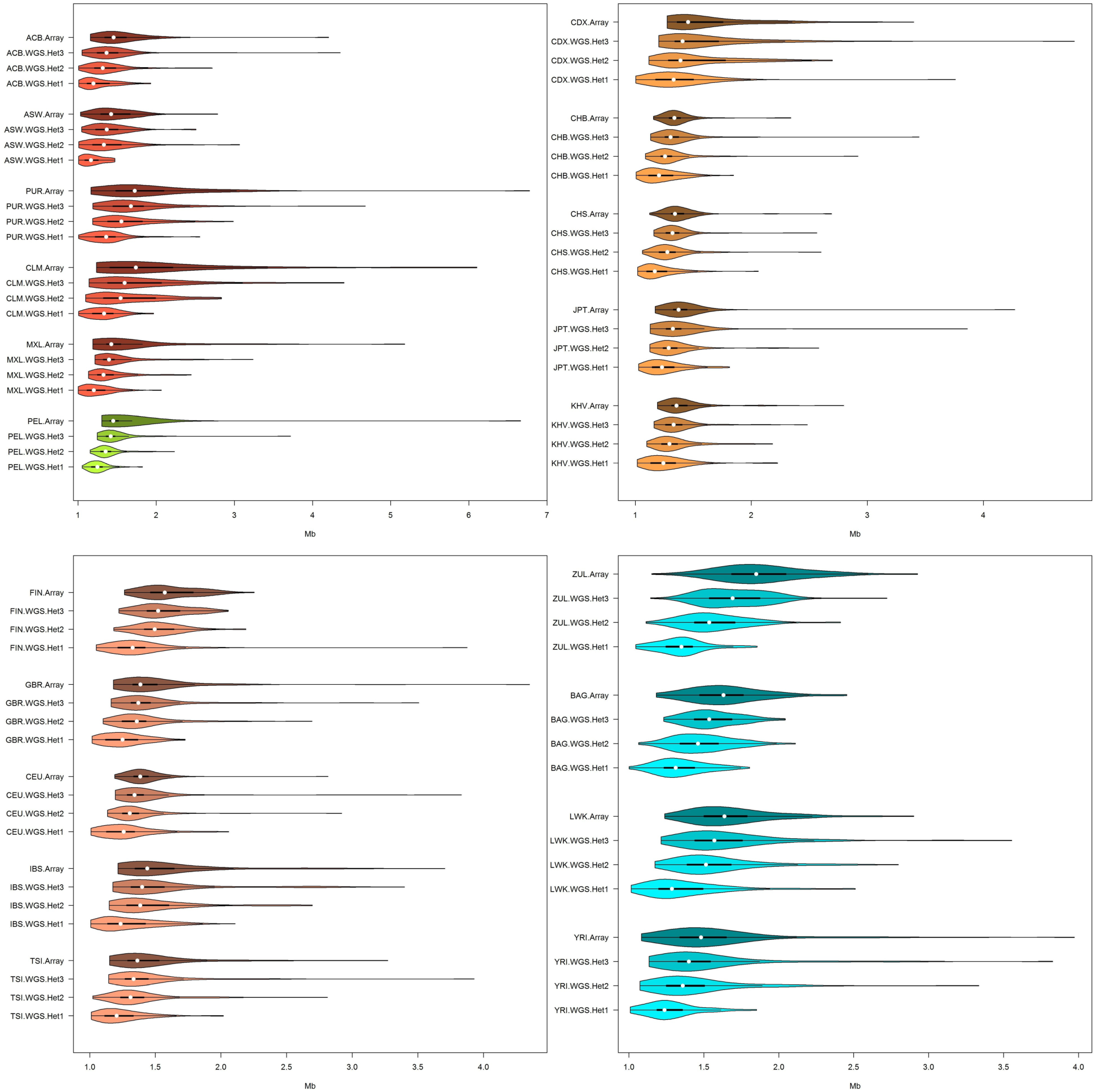
Violin plots of mean ROH size longer than 1 Mb (in Mb). Different biogeographical groups have different *x*-axis scales in an attempt to maximize the difference between distributions within populations. See Figure 2 legend for population codes.

**Figure 4.**
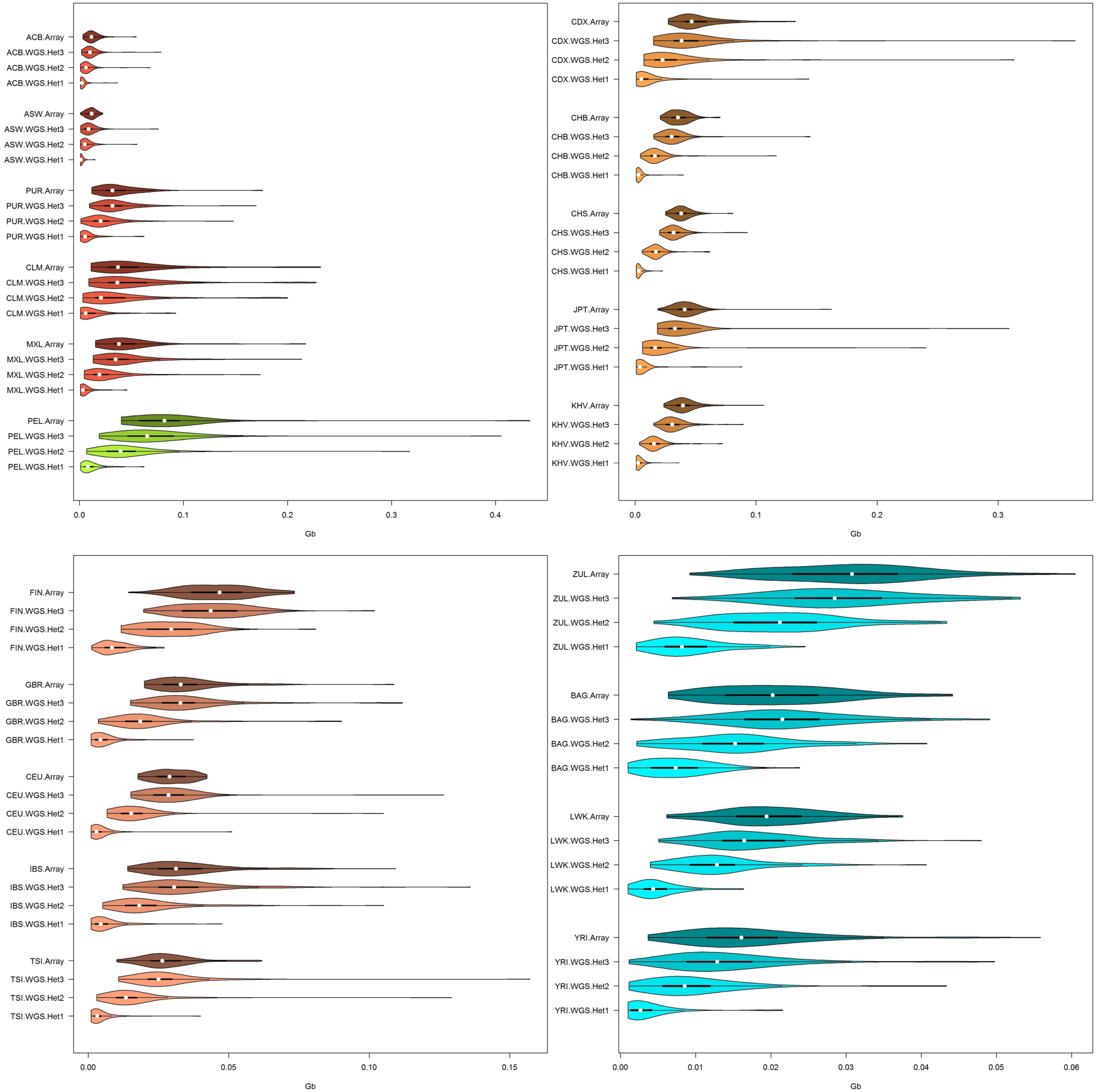
Violin plots of mean total sum of ROH longer than 1 Mb (in Gb). See figure 2 legend for population codes.

**Figure 5.**
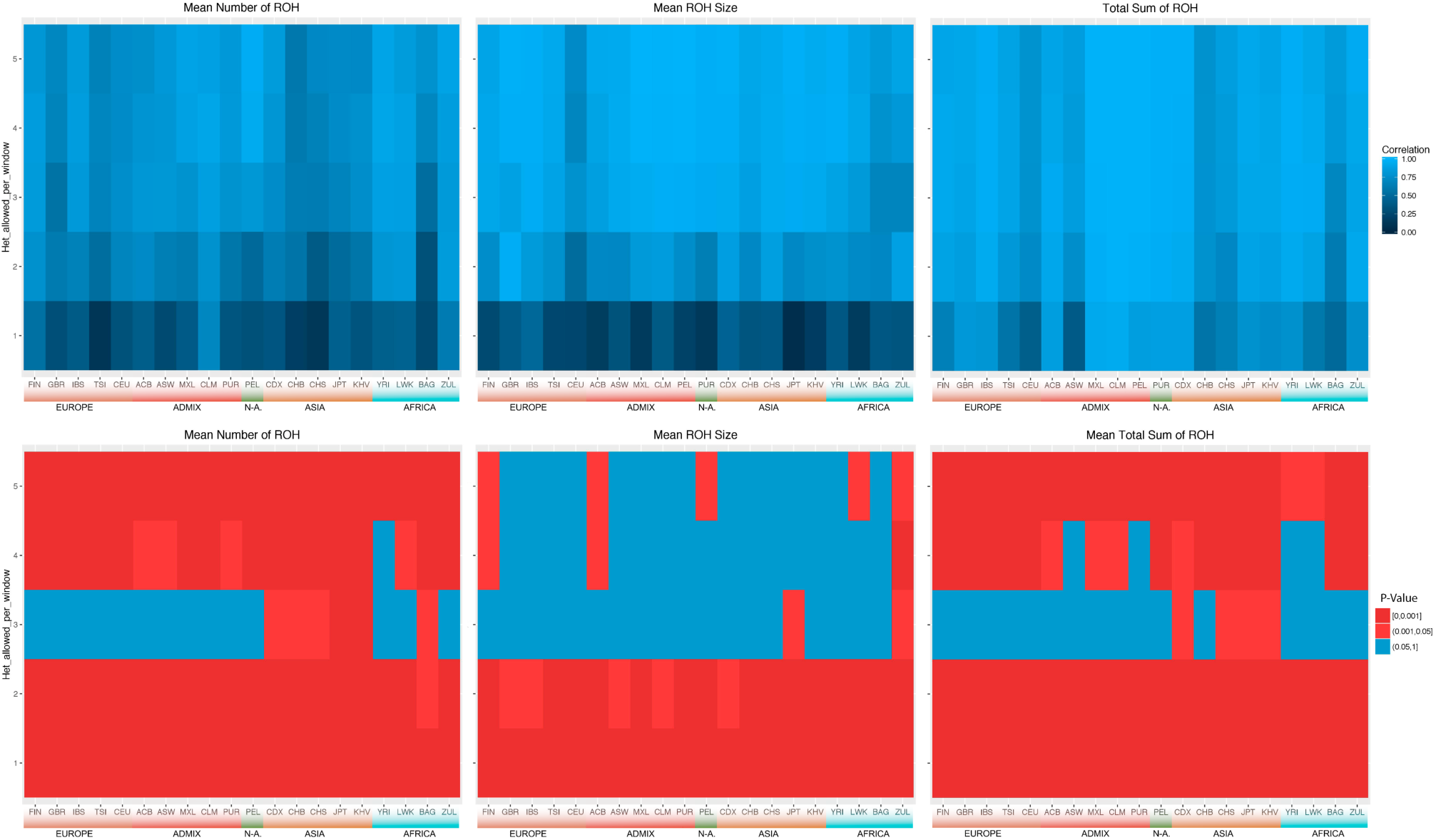
Heatmaps of correlations and MWW tests of mean number of ROH, mean ROH size and mean total sum of ROH between array data allowing 1 heterozygous SNP per ROH and WGS data allowing 1 to 5 heterozygous SNPs per ROH (y-axis). **Top row of images|** Pearson correlations. **Bottom row of images|** P-values of Mann-Whitney-Wilcoxon non-parametrical test (MWW), red shows significant difference between array and WGS while blue shows distributions that cannot be considered different. See Figure 2 legend for population codes.

### Comparing ROH with different lengths

Considering WGS data with 3 heterozygous SNPs allowed per window, the best PLINK condition to obtain ROHs comparable with ones obtained from array data, it is interesting to compare the mean sum of ROH, between technologies, in different ROH length categories (Figure 6). This is relevant because the study of different ROH lengths has different applications, as indicated in Table 2. Figure 6 shows that for ROH longer than 1Mb, the array and WGS mean total lengths are very similar, with some exceptions like the JPT, in the case of ROH longer than 8 Mb. However, WGS data systematically detected more short ROH (0.3 – 1Mb) than array data. This outcome is expected and is caused by the lower SNP coverage of array data, since PLINK considered just ROH containing at least 50 SNPs. This gap between array and WGS data can be corrected for small ROH by changing PLINK parameters and relaxing the number of SNPs needed to call a ROH (--homozyg-snp 30, data not shown).

**Figure 6.**
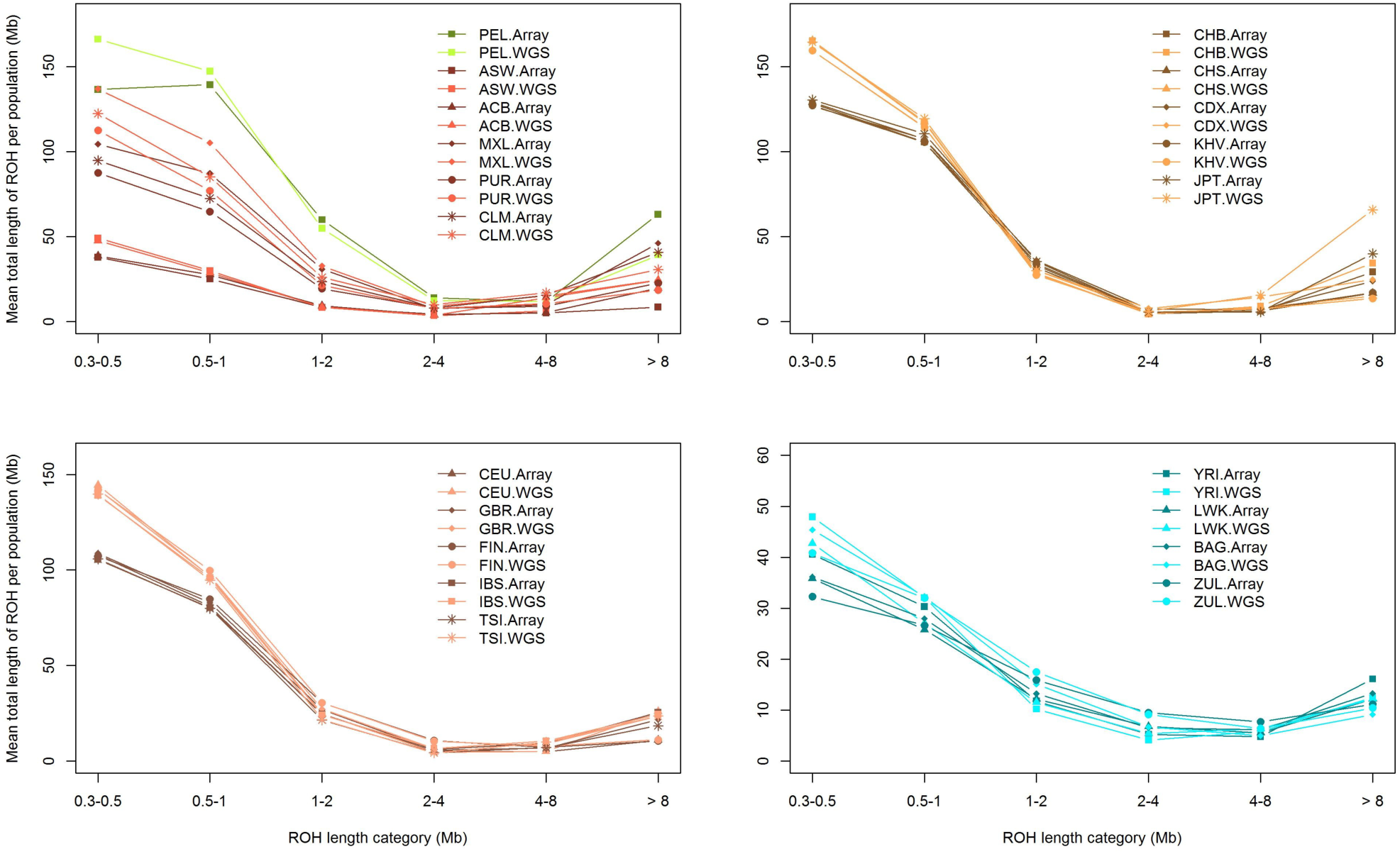
Mean sum of ROH in different length categories. The light colored lines represent WGS with 3 heterozygous SNP allowed per window and dark colored lines represent array data with 1 heterozygous SNP allowed per ROH. See Figure 2 legend for population codes.

**Table 2.** Performance of different technologies (array, WGS low coverage and WES) with different ROH size classes (Short < 1Mb, Medium 1 – 8Mb and Long > 8Mb).

| ROH size Class | SNP Array | WGS low coverage | WES | Applications |
| --- | --- | --- | --- | --- |
| Short < 1Mb | Poor performance due to low SNP coverage. Can be adjusted to detect ROH by modifying the number of SNPs required in a ROH. | Able to detect but need to build adjustment for genotype calling errors. | Able to detect but only in selected genomic regions. Software like H<sup>3</sup>M<sup>3</sup> allows meaningful regional analysis. | Detection of rare variants involved in deleterious recessive alleles and directional dominance. Analysis of LD patterns and extreme bottle necks. |
| Medium 1-8Mb | Able to detect if the array has at least 300K SNPs. ROH boundaries will be fuzzier in comparison with WGS low coverage data. | Good performance but need to build adjustment for genotype calling errors. Allowing 3 heterozygous SNPs per ROH would grant meaningful outcomes. | Able to detect, but only in selected genomic regions and boundaries of ROH could be fuzzy if they reach into non-exonic regions. | Detection of rare variants involved in diseases. Analysis of inbreeding depression. Genome architecture and ROH island detection. Population history, bottle necks, remote consanguinity and genetic drift. |
| Long > 8Mb | Good performance if the array has at least 300K SNPs. | Good performance but need to build adjustment for genotype calling errors. Allowing 3 heterozygous SNPs per ROH would grant meaningful outcomes. | Poor performance due to short size of most exons and their sparsity across the genome. | Analysis of inbreeding depression. Validation of GWAS findings. Population history and cultural practices, close consanguinity. |

## Discussion and Conclusions

Runs of Homozygosity were first detected due to the ability to perform denser genome-wide microsatellite marker scans in the late 1990s (Broman et al. 1999). Soon after the first ROH study using short tandem repeat polymorphisms (STRPs) was released, the first SNP arrays started to become available. During those first years, using arrays with densities of 40K and 120K SNPs, ROH were discovered to be ubiquitous across all human populations (Gibson et al. 2006). However, it was not until the first arrays with more than 300K SNPs were used that the analysis of ROH started to shed some light onto the understanding of human demographic history and in deciphering the genetic structure of traits and complex diseases (Ku et al. 2011; McQuillan et al. 2012; Wang et al. 2015). Currently array-based genotyping covers around 1.9 to 2.2 million SNPs, allowing meaningful detection of ROH longer than 1Mb, and even though this is an important improvement over previous arrays, it covers only ∼2% of the total common SNPs present in the human genome (LaFramboise 2009; Lamy et al. 2011). This prevents the use of array data for detecting shorter ROH, an essential component contributing to the understanding of human genetics. WGS will soon allow shorter ROH to be more reliably called; permitting the effect of very short ROH on diseases risk to be quantified. Thus, analyzing the effect of different lengths of ROH may reveal the relative contributions of rare and common variants to the demographic history of human populations.

Ideally, WGS deep coverage would be the best option to study ROH, since genotype calling will be robust for low MAFs and ROHs of virtually any size would be detected. However, two major issues prevent the use this technology. First, the lack of WGS deep coverage data for population studies and secondly, the extreme computational expense of analyzing this type of data using current software. Unlike deep coverage, low coverage WGS data is more abundant and affordable, and the computational effort of obtaining ROH is less computationally intensive. The only drawback of using this data is the calling error associated with it. By comparing ROH obtained from array data, we demonstrate in this article that this problem can be mitigated by allowing 3 heterozygous SNPs per window using PLINK software to obtain ROH longer than 1Mb. In all populations, the highest correlation was achieved when allowing 3 to 4 heterozygous SNPs per window (Figure. 5 top row of images). Regarding MWW tests (Figure. 5 bottom row of images), unlike mean number and total sum of ROH, for most of the populations, mean ROH size remains equivalent between technologies when allowing 3 or more heterozygous SNPs per window. As expected, we get more ROH by allowing more heterozygous SNPs, but the mean size remains constant. As a consequence, the mean total sum of ROH increases with more heterozygous SNP allowed.

Interestingly, four populations from East and South Asia did not conform to the patterns observed in the other populations; in fact for the Dai and Han populations from China (CDX, CHS), Kinh population from Vietnam (KHV) and the Japanese population (JPT), it was not possible to obtain the same mean number and total sum of ROH between array and WGS data. This may be explained by population structure, but perhaps the inferior performance of the Infinium Omni 2.5-8 Bead chip in Asian populations (Ha et al. 2014) is the more plausible explanation. This could also explain why it was not possible to obtain same number of ROH in the Baganda population from Uganda (BAG) or the same mean ROH size in the Zulu population from South Africa (ZUL).

WGS data present the ability to identify shorter ROH (Figure 6), however it would be important to compare the short ROH detected using low coverage, compared to high coverage data to establish a comparative analysis guideline. In Table 2 we present a comparison in performance of the application of three different technologies (SNP array, WGS low coverage and WES data) to detect short, medium and long ROH.

This study provides evidence-based guidelines for the combined analysis of array and low coverage WGS data when studying ROH to investigate population history and to detect associations with complex diseases and traits. To date, ROH studies remain underexplored in the search for genetic variants associated with common diseases in different populations and in the detection of signals of selection.

## Materials and Methods

### Description of Data

Individuals with both genome-wide SNP genotypic data and WGS low coverage data from the 1000 Genomes Project – Phase 3 (KGP) (Sudmant et al. 2015; The 1000 Genomes Project 2015) and the African Genome Variation Project (AGVP) (Gurdasani et al. 2015) were used. For both datasets the Infinium Omni 2.5-8 Bead chip from Illumina was used. The KGP, includes a total of 1685 individuals from 18 populations with genotypic data available from array and WGS low coverage (4x). From Europe: FIN (Finish in Finland, n=99), GBR (British in England and Scotland, n=91), IBS (Iberian populations in Spain, n=105), TSI (Tuscany in Italy, n=102) and CEU (Utah residents with European ancestry=99). From America: ASW (Americans of African ancestry in Huston, n=61), ACB (African Caribbean in Barbados, n=96), PUR (Puerto Rican in Puerto Rico with admix ancestry, n=104), PEL (Peruvian in Lima, Peru with Amerindian ancestry, n=85), CLM (Colombian in Medellin, Colombia with admix ancestry, n=95) and MXL (Mexican with admix ancestry in Los Angles, USA, n=100). From Asia: CDX (Chinese Han in Xishuangbanna, China, n=98), CHB (Chinese Han in Beijing, China, n=100), CHS (Southern Han Chinese, n=105), JPT (Japanese in Tokio, Japan, n=100) and KHV (Kinh in Ho Chi Minh city, Vietnam n=99). From Africa: YRI (Yoruba in Ibadan, Nigeria, n=108) and LWK (Luhya in Webuye, Kenya, n=99). The AVGP includes 2185 samples from 16 African populations; we use WGS data for two: 100 Zulu from South Africa and 100 Baganda from Uganda, where genotype data from the Omni 2.5-8 SNP array and WGS data at 4x coverage are available. Only SNPs of the 22 autosomes were included in this analysis. For each population, data from both array genotyping and WGS were filtered to remove SNP with minor allele frequencies lower than 0.05 and those that divert from H-W proportions with *p* < 0.001. This filtering limits the effects of ascertainment bias caused by the small number of individuals in the SNP discovery panel, in the case of the array, and the calling errors associated with a low depth coverage of whole genome sequence data.

### Identification and Characterization of ROH

We use PLINK v1.9 to identify ROH. The following conditions where used:

-- *homozyg-snp 50.* Minimum number of SNPs that a ROH is required to have
-- *homozyg-kb 300.* Length in Kb of the sliding window
-- *hmozyg-density 50.* Required minimum density to consider a ROH (1 SNP in 50Kb)
-- *homozyg-gap 1000.* Length in Kb between two SNPs in order to be considered in two different segments.
-- *homozyg-window-snp 50.* Number of SNPs that the sliding window must have
-- *homozyg-window-het (1 to 5).* Number of heterozygous SNP allowed in a window
-- *homozyg-window-missing 5.* Number of missing calls allowed in a window
-- *homozyg-window-threshold 0.05.* Proportion of overlapping windows that must be called homozygous to define a given SNP as in a “homozygous” segment.

The minimum length of a ROH was set to 300 kb. PLINK allows the setting of different variable number of heterozygous SNPs per window, with a default value of 1 heterozygous genotype per ROH, in order to tolerate genotyping calling errors (*--homozyg-window-het 1*). This is especially relevant in dealing with WGS low coverage data and therefore we are testing the equivalence between ROH obtained from array genotyping and WGS data.

Our goal is to determine under which conditions detecting ROHs using low coverage sequence data results in the comparable results as using array data. There are several characteristics of ROHs we can measure and we would like these characteristics to be the same no matter the technology used.

Our first characteristics is *ep(P,h)*, a measure of the empirically observed number of heterozygous SNPs found in ROHs in population P when we allow h heterozygous SNPs (see supplementary material for the definition). This observed number of heterozygous SNPs differs from the parameter used for detecting ROHs depending on the population and technology platform characteristics. Figure 1 shows *ep(P,h)* for different populations. *ep(P,h)* values for different populations (*P*) and heterozygous SNP allowed (*h*) are shown in Supplemental_Table_S1. These results show that for low coverage sequence data, *h* should be at least 3 to get the same characteristics as array data. The analysis below explores this in more detail.

### Statistical Analysis

For comparison purposes three variables per population were defined. Mean number of ROH as the mean number of ROH longer than 1Mb. Mean ROH size as the mean size of ROH longer than 1Mb. Total sum of ROH as the mean total sum of ROH longer than 1Mb. Considering just ROH longer than 1Mb allows the selection of only the ROH arising from identity by descent and to remove any LD effects. Data distributions were illustrated using violin plots. This plot combines a box plot with a kernel density plot, where the interval width is obtained by the rule of thumb. The violin shows a colored kernel density trace with the interquartile range as a black line and median as a white dot. This representation is especially relevant when dealing with data or variables that show skewed distributions and is a good means of comparison between populations, when dealing with asymmetric distributions where the median is more informative than the mean. Statistical comparisons between mean number of ROH, mean ROH size and mean total sum of ROH for different populations, technologies and PLINK conditions were performed by Pearson’s correlation and Mann-Whitney-Wilcoxon non-parametric test (MWW). All the exploratory and statistical analyses were performed using R.

## Data access

African Genome Variation Project data (AGVP) are available from the European Genome-phenome Archive (EGA, http://www.ebi.ac.uk/ega/), hosted by the EBI, under accession Numbers EGAS00001000363 and EGAS00001000286.

## Acknowledgements

FCC is a National Research Foundation of South Africa (NRF) postdoctoral fellow and MR holds a South African Research Chair in Genomics and Bioinformatics of African populations hosted by the University of the Witwatersrand, funded by the Department of Science and Technology and administered by the NRF. This work was enabled through a 6-month scientific exchange visit of FCC to the group of Jim Wilson at the University Edinburgh (supported by funding allocated by Nick Hastie (UK) and MR (SA)).

